# Effects of microtubule length and crowding on active microtubule network organization

**DOI:** 10.1101/2022.09.23.509184

**Authors:** Wei-Xiang Chew, Gil Henkin, François Nédélec, Thomas Surrey

## Abstract

Active filament networks can organize into various dynamic architectures driven by crosslinking motors. Densities and kinetic properties of motors and microtubules have been shown previously to determine active microtubule network self-organization, but the effects of other control parameters are less understood. Using computer simulations, we study here how microtubule lengths and crowding effects determine active network architecture and dynamics. We find that attractive interaction mimicking crowding effects or long microtubules both promote the formation of nematic networks of extensile bundles instead of contractile networks. When microtubules are very long and the network is highly percolated, a new isotropically motile network state resembling a ‘crawling mesh’ is predicted. Using *in vitro* reconstitutions, we confirm the existence of this crawling mesh experimentally. These results provide a better understanding of how active microtubule network organization can be controlled, with implications for cell biology and active materials in general.

## INTRODUCTION

Active filament networks, driven out of equilibrium by ATP-consuming crosslinking motor proteins, have the capacity to adopt different dynamic organizations. In living cells, they play important roles in various processes such as spindle assembly (Kapoor 2017), cytoplasmic streaming (Quinlan 2016), and cell shape control (Salbreux, Charras and Paluch 2012). Biochemical reconstitutions with purified proteins *in vitro* have made important contributions to our understanding of active network organization and dynamics (Needleman and Dogic 2017, Koenderink and Paluch 2018, Dogterom and Surrey 2013, Alfaro-Aco and Petry 2015). Networks formed from microtubules and motors can display contractile (Nedelec et al. 1997, Foster et al. 2015, Hentrich and Surrey 2010, Surrey et al. 2001, Torisawa et al. 2016) or nematic, extensile behavior (Sanchez et al. 2012, Lemma et al. 2022, Roostalu et al. 2018, Henkin et al. 2022). Contraction and expansion are fundamentally opposite activities and the molecular parameters that determine these collective behaviors remain incompletely understood.

Microtubules are dynamic, structurally polar filaments with distinct ‘plus’ and ‘minus’ ends, composed of tubulin subunits arranged into a tube. Early *in vitro* experiments with artificially oligomerized plus-end directed kinesin-1 and microtubules growing in solution showed the formation of contractile networks, eventually forming asters with a radially polar microtubule arrangement (Nedelec et al. 1997, Surrey et al. 2001). Computer simulations demonstrated that aster formation depends on the motor’s ability to remain bound upon reaching microtubule ends, allowing the ends to be brought together. Similar contractile networks were later observed in the presence of the natural microtubule crosslinking motors kinesin-5 and kinesin-14, which walk towards the plus or minus ends, respectively, and form asters with opposite polarity (Hentrich and Surrey 2010, Roostalu et al. 2018). Actin filaments and related motors also readily form contractile networks *in vitro* (Alvarado et al. 2017, Koenderink and Paluch 2018) and contractility often drives the function of actin networks *in vivo*, e.g. the actomyosin cytokinetic ring or the cell cortex, (Murrell et al. 2015, Levayer and Lecuit 2012), due to the lower filament stiffness (Stam et al. 2017). In contrast, microtubule networks more often rely on their ability to expand, as illustrated by the mitotic spindle, where the separation of the chromosomes often depends on anaphase expansion.

Extensile nematic microtubule network formation was first reconstituted *in vitro* using short pre-polymerized microtubules and artificially oligomerized kinesin-1 motors, but now in the additional presence of a crowding agent causing microtubule bundling by depletion forces (Sanchez et al. 2012). With fewer crowding agents, or a reduced density of static microtubules, the networks were shown to become contractile (Lemma et al. 2022). Later it was shown that microtubules growing in solution in the absence of a crowding agent can also be organized only by motors into extensile nematic networks, provided the tubulin concentration is high enough to promote fast microtubule growth and high microtubule densities (Roostalu et al. 2018). Computer simulations showed that a higher motor speed, compared to the microtubule growth speed, facilitated the accumulation of the motor at microtubule ends, causing network contraction and aster formation, whereas relatively fast microtubule growth and high microtubule densities favor extensile nematic network formation (Roostalu et al. 2018).

Altogether, qualitatively similar networks have therefore been observed both with and without crowding agents, and with both shorter and longer microtubules. While theoretical models and computer simulations have helped to explain the effects of certain control parameters in self-organized microtubule-motor networks (i.e. microtubule growth rate, microtubule density, motor speed, motor density, motor composition) (Blackwell et al. 2016, Belmonte, Leptin and Nedelec 2017, Roostalu et al. 2018, Fürthauer, Needleman and Shelley 2021) other parameters that appeared determinant in *in vitro* studies remained unexplored. Particularly, a theoretical exploration of the effects of crowding induced depletion forces and microtubule length has not been performed yet, limiting our consolidated understanding of active microtubule network organization.

Here we explore the effects of these system parameters on simulated active networks composed of microtubules and motors. We find that short-range attractive forces between microtubules, mimicking depletion forces, promote bundling, thereby preventing aster formation, and particularly at high microtubule densities generate networks of extensile bundles. In the absence of such short-range attractive forces, shorter microtubules promote the formation of contractile active networks, whereas long microtubules promote nematic networks of extensile bundles, or when the degree of percolation is high a isotropic motile network resembling a ‘crawling mesh’, a new network state whose existence we also demonstrate here experimentally. The control parameters determine network organization by affecting the numbers of motors that accumulate near the end of microtubules, or on their side, and in these different configurations, the motors can tip the outcome of active microtubule network organization towards contractile or extensile states.

## RESULTS

We simulated active networks consisting of microtubules and microtubule-crosslinking motors using Cytosim. Microtubules and motors were modelled essentially as described earlier (Nedelec 2007, Henkin et al. 2022) (see Fig. 1A and Methods). Microtubules grew in a thin and flat three-dimensional geometry from a fixed number of nucleators by plus-end elongation and repelled each other via soft-core interactions. Motors with the ability to bind two different microtubules could organize them into active networks. In this work, the motor properties mimicked those of the human spindle motor KIF11, a plus-end directed symmetric motor.

**Figure 1.**
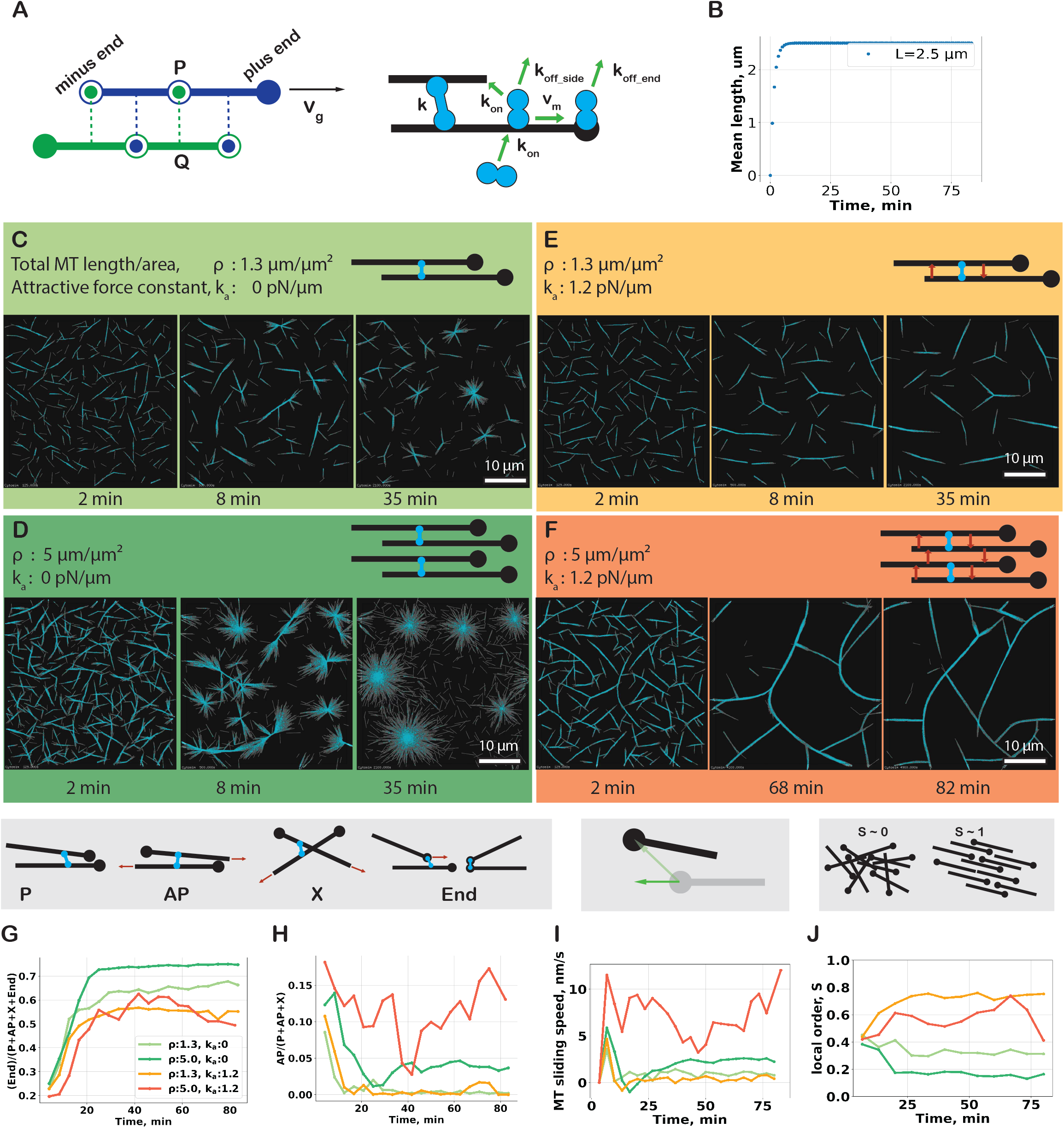
(A) Elements of the microtubule/motor simulation. (Left) Microtubule filaments are discretized into points separated by equal distance, allowing a filament to bend but not to stretch. Each point is subjected to forces from bending elasticity, and interaction with crosslinker and other filaments. For the steric interaction, we consider all constitutive points P of a filament and project on the segments of other filaments. The force is orthogonal to the opposite filament at the projection point Q. An opposite force is applied to the first filament in P. The attractive force is implemented in the same way. Microtubules only grow at the plus end with a constant rate. (Right) Motors have two microtubule-binding units. A free motor unit binds to a filament with a binding rate k_on_. Motors that connect two filaments form a Hookean crosslink. A bound motor unit moves with a speed that is linearly dependent on the force, as defined by the stall force and the unloaded speed v_m_. A motor can detach with a rate k_off_side_ or k_off_end_ depending on its position along the filament. (B) Time course of the microtubules’ mean length. Time course of microtubule (gray) organization at (C) low total microtubule length per area (1.3 μm/μm^2^) and (D) high total length per area (5 μm/μm^2^) in the presence of KIF11 motor (cyan) without attractive interfilament depletion force. Time course of active microtubule organization in the presence of attractive depletion force ka (1.2 pN/μm) at (E) low and high (F) total microtubule length per area. Motor crosslinks are categorized depending on the angle between the microtubules into P, AP or X links, and whether they occur near the microtubule minus ends. P links connect parallel microtubules where the internal angle is smaller than 60°. AP links connect antiparallel microtubules at an angle between 120° and 180°. X links connect microtubule sides when these microtubules form an angle from 60° to 120°. End-links connect one or both microtubule ends. Time series of (G) the fraction of endbound motors to all types of crosslinks, (End links)/(P+AP+X+End) (see Methods), (H) the fraction of side-bound motors that forms antiparallel links, AP/(P+AP+X), (I) the mobility of microtubule minus ends along the filament axis, and (J) the local nematic order parameter calculated with a sampling window size of 10 μm. The KIF11 motor-to-microtubule ratio is 16. The simulation extends for 80 min in a box with dimensions 40 μm × 40 μm × 0.2 μm with periodic boundary conditions. See also Movie 1.

### Effects of a short-range attractive force between microtubules on active network organization

We began by studying the effect of a short-range attractive force between microtubules, in systems with different microtubule densities. These forces promote the formation of bundles in which adjacent microtubules are free to slide longitudinally relative to each other, and thus mimic depletion forces induced by crowding agents in experimental active networks (Sanchez et al. 2012). Microtubules grew with a speed that was initially equal to the motor speed, then growth slowed down and finally stopped after ~ 8 min when microtubules reached an average length of 2.5 μm (Fig. 1B). Motors remained bound at microtubule ends for an average of 5 s, allowing them to form asters within 35 min in the absence of an attractive depletion force (Fig. 1C, D), as observed previously in experiments and simulations (Roostalu et al. 2018, Henkin et al. 2022). Whereas microtubules contract locally into small disconnected asters at a lower microtubule density (Fig. 1C), at a higher microtubule density the network initially percolated but subsequently broke down, finally contracting into larger asters (Fig. 1D), as indicated by the large number of microtubule end-bound motors (Fig. 1G).

Under these conditions, introducing an attractive force between the microtubules suppressed aster formation, instead causing the formation of microtubule bundles (Fig. 1E, F), similar to experiments with a crowding agent (Lemma et al. 2022). Isolated parallel microtubule bundles or parallel bundles connected by their plus ends formed at a lower microtubule density (Fig. 1E) in which individual microtubules were relatively static, as indicated by a slow average microtubule sliding speed (Fig. 1I) and a small number of motor links connecting antiparallel microtubules (Fig. 1H). At a higher microtubule density, bundles extended and collapsed onto each other leading to turbulent network activity (Fig. 1F) (Movie 1), displaying fast microtubule sliding with a fluctuating speed (Fig. 1I) and a large, but also fluctuating number of motor links between antiparallel microtubules (Fig. 1H). The turbulent behavior of the extensile bundles was also reflected by a relatively high, fluctuating local nematic order parameter (Fig. 1J). The turbulent network behavior of extending, bending and recombining bundles in these simulations is remarkably similar to what has been observed in experimental active networks in the presence of crowding agents (Sanchez et al. 2012).

Next, we explored a larger part of the organizational phase space to elucidate more systematically the combined effects of varying both the strength of the short-range attractive force and the microtubule density (Fig. 2A). Extracting the local nematic order parameter from the simulated end states showed that generally, increasing the attractive force leads to more local nematic order (Fig. 2B). The lowest degree of local nematic order was observed when almost all microtubules were incorporated into asters, whereas bundling increased the order parameter. For the highest attraction forces and the densest systems the average microtubule sliding speed was highest (Fig. 2C), correlating with the largest fraction of motor crosslinks engaged in antiparallel microtubule contacts (Fig. 3A ii), demonstrating that microtubules fail to polarity sort in the turbulent regime. The measured nematic order was lower, as a consequence of a temporal fluctuation due to turbulent network motion.

**Figure 2.**
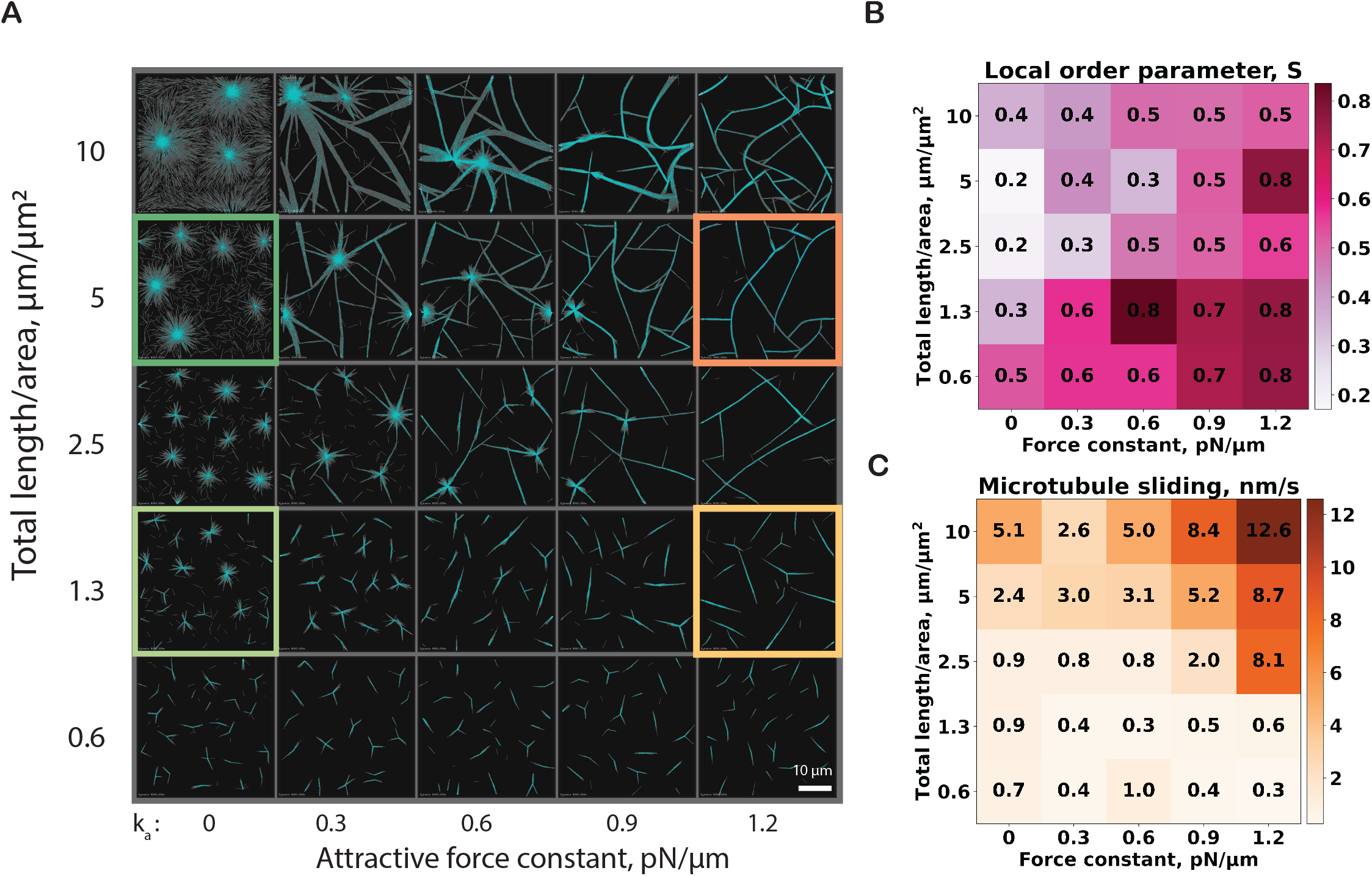
(A) Organizational phase space of the microtubule network at various combinations of total microtubule length per area and attractive force strength. The four colored squares correspond to conditions in Fig. 1 C-F. All boxes are the same size. (B) Local nematic order measured for each simulation shown in (A) at 80 min. (C) Calculated mobility of microtubules for each simulation shown in (A). For B and C, the color scales are linear as indicated (right). The ranges correspond to the minimum and maximum values in the entire set.

**Figure 3.**
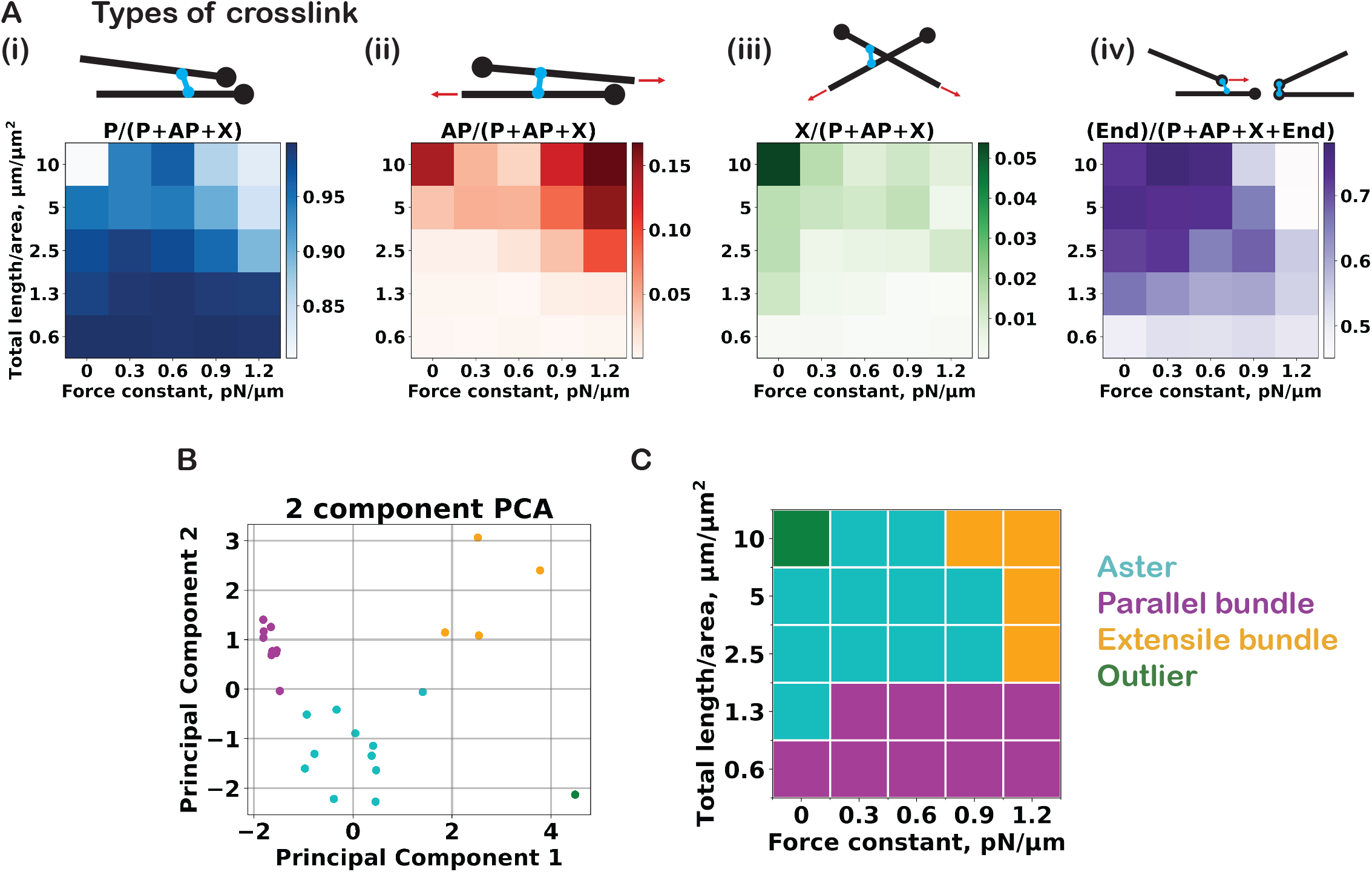
(A) The fraction of each link type is represented using color shades for different simulations, varying microtubule density and attractive force strength. (i) P links, (ii) AP links, (iii) X links and (iv) end links. The color scales are linear, corresponding to the minimum and maximum observed in the entire dataset. Original numerical values of the same data are reported in Fig. S3A. (B) Scatter plot of the 25 microtubule organizations in the subspace of the two greatest principal components. The four clusters identified via the K-means analysis are indicated by different colors: cyan as aster, magenta as parallel bundle, orange as extensile bundles and green as outlier. (C) The corresponding regimes in the density-attractive force constant plot.

We then categorized the microtubule organizations based on six network descriptors: the scalar nematic order parameter (Fig. 2B), microtubule speed (Fig. 2C), parallel links (Fig. 3Ai), antiparallel links (Fig. 3Aii), X crosslinks (Fig. 3Aiii) and end links (Fig. 3Aiv), dividing the phase space into three regimes. A clustering procedure was performed in the subspace of the two main principal components of the descriptors in an unsupervised manner (see Method) (Fig. 3B, Fig S1A). The identified clusters (Fig. 3C) correspond well to (i) the contractile aster-forming regime, characterized by low order parameter and a large fraction of end links (Fig. S1B), (ii) the parallel bundle state, with a high nematic order parameter and a large fraction of parallel links, and (iii) turbulent networks of extensile bundles, with high microtubule sliding speed and a large fraction of antiparallel links. Note that the organization at the highest density without the attractive force that appears to be a focused aster with radially aligned microtubules is an outlier that shares characteristics of (i) and (iii) as can be seen in the principal component plot (bottom right of Fig. 3B).

Together, these simulations show that both increasing the attractive force between microtubules and the microtubule density promotes a turbulent network of extensile microtubule bundles.

### Effects of microtubule length on active network organization

Next, we studied the effect of the microtubule length on active microtubule organization. This was done in the absence of a short-range attractive force between microtubules to mimic *in vitro* experiments in which the transition between contractile and nematic was studied by varying the tubulin concentration in the absence of crowding agents (Henkin et al. 2022). In our simulations, we systematically varied the maximum length of the microtubules and the microtubule density, defined as the total length of all microtubules per area (Fig. S2B). Increasing the microtubule length from 2.5 μm to 10 μm at an intermediate microtubule density (2.5 μm total microtubule length per μm^2^) prevented individual aster formation and led to the formation of a polarity-sorted network containing polar bundles (Fig. 4A, Fig S2A, Movie 2). Concomitantly, the local order parameter increased from 0.2 to 0.9 (Fig 4B) and the average microtubule sliding speed increased from 1 nm/s to 6 nm/s (Fig. 4C). Further increasing the microtubule length caused a slight reduction of the nematic order to 0.7 but a continued increase of the microtubule sliding speed up to 17 nm/s for the longest microtubules.

**Figure 4.**
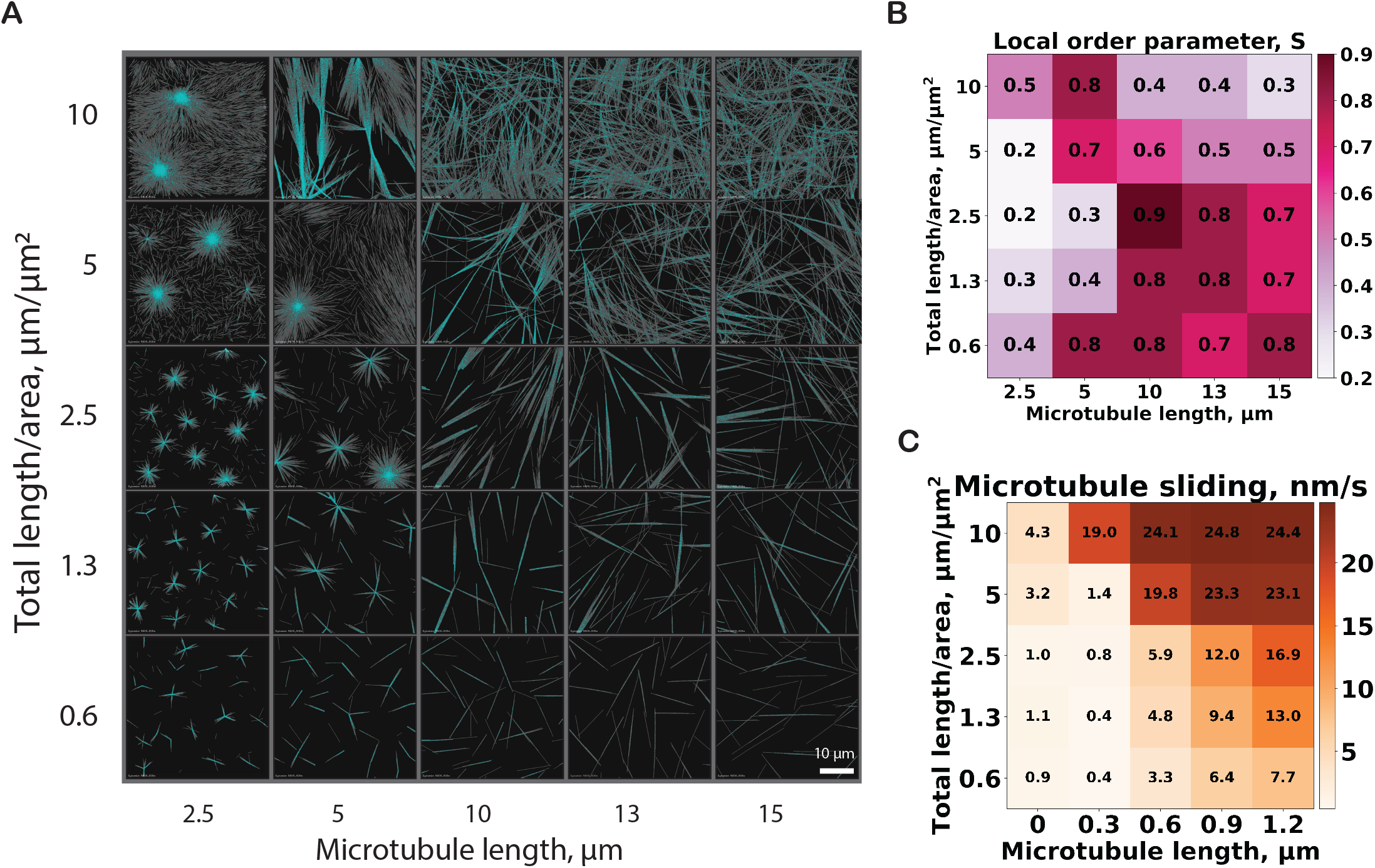
(A) Organization of microtubule networks depending on average microtubule length and microtubule density. The KIF11 motor-to-microtubule ratio is 16. The simulation was performed for 60 minutes in a box of dimensions Lx μm × Ly μm × 0.2 μm, where Lx = Ly = 16 × (microtubule length). For visualization purposes, only a part of the entire simulation space is shown with a constant area of 40 μm × 40 μm. The entire simulation spaces are shown in Fig. S2. (B) Local nematic order measured in each simulation shown in (A) at 60 min. (C) Calculated mobility of microtubule minus ends for all simulations shown in (A). For B and C, the color scales are linear, corresponding to the minimum and maximum values observed in the dataset. See also Movie 2, 3.

At higher microtubule densities (10 μm total microtubule length per μm^2^), the same trend was observed for simulations with increasing microtubule length (Fig. 4A, Movie 3A). However, the network with the maximum order parameter, consisting of extensile bundles, was reached already at the shorter microtubule length of 5 μm, followed by a decrease in the order parameter with further increases in microtubule length (Fig. 4B). The microtubule sliding speed again increased continually with increasing microtubule length approaching 25 nm/s (close to the speed of the motors at 30 nm/s) for the longest microtubules (Fig. 4C).

Clustering analysis identified four distinct groups in the microtubule length/density phase space (Fig 5B, Fig 5C). Contractile networks forming asters were generated when microtubules were short. This regime was characterized by low nematic order (Fig. S1D) and slow average microtubule motility. Parallel bundles formed when microtubules were long at low to intermediate microtubule densities. They were characterized by high nematic order and very slow microtubule motility. The fastest microtubule sliding was observed for the longest microtubules and highest microtubule densities. Here the order parameter was rather low, indicative of a mostly isotropic, but highly motile network. In between these regimes was the regime of extensile bundles that was characterized by the high values for the order parameter, but lower sliding speeds than for the isotropic motile network.

**Figure 5.**
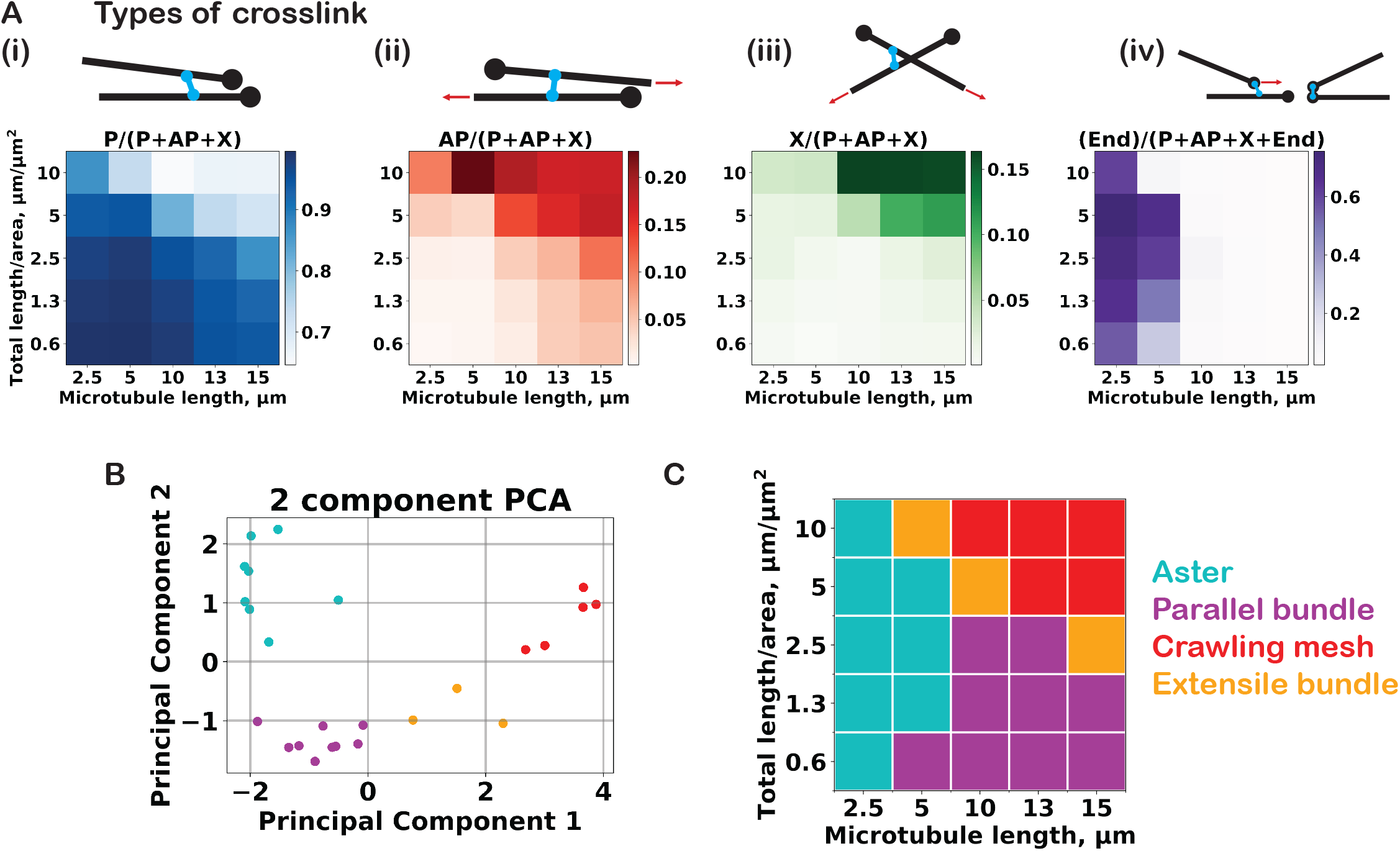
(A) Fraction of the different category of motor crosslinks at each microtubule density and attractive force strength: (i) P links, (ii) AP links, (iii) X links, and (iv) end links. The color scales are linear, corresponding to the minimum and maximum values in the entire dataset. Numerical values are reported in Fig. S3A. (B) Scatter plot of the 25 microtubule organizations in the subspace of the two greatest principal components. The four clusters identified via the K-means analysis are: aster (cyan), parallel bundle (magenta), extensile bundles (orange), and crawling mesh (red). (C) Classification indicated in the density-length plot.

The formation of these four distinct network architectures can be explained by investigating the types of crosslinks that the motors form. In the aster regime motors have efficiently accumulated at microtubule ends, as indicated by a high fraction of end links, promoted by microtubules being short (Fig. 5A iv, Fig S1D). In the regime of extensile bundles, one can observe a maximum of motors crosslinking antiparallel microtubules, promoted by the ability of the motors to align microtubules of intermediate length and density (Fig. 5A ii). For the longest microtubules and the highest densities, the networks become highly percolated with many crosslinks between non-aligned microtubules (X links in Fig. 5A iii). This network remains fairly isotropic, because the high degree of percolation hinders microtubule alignment or end gathering. Instead, microtubules translocate through the network roughly following the direction of their axis (Movie 3B), a previously not described state which is isotropically motile since microtubules move in all directions equally, giving the impression of a ‘crawling mesh’.

As the contractility or expansion of the network is correlated with the number of end-links, the organizational state is expected to be dependent on the motor’s end-unbinding rate, a previously established control parameter for motor-microtubule networks (Surrey et al. 2001, Tan et al. 2018) which modulates the relative fraction of end-versus side-bound motors. We should therefore be able to change the location of the boundary between contractile and expansile states by adjusting this rate. We find that the network states characterized by high microtubule sliding speeds indeed occupy a larger area of the parameter space when the end unbinding rate is increased (Fig. 6A, B), corresponding to decreased end-accumulation (Fig. 6C), leading to suppression of aster formation and therefore promotion of extensile bundling (Movie 4).

**Figure 6.**
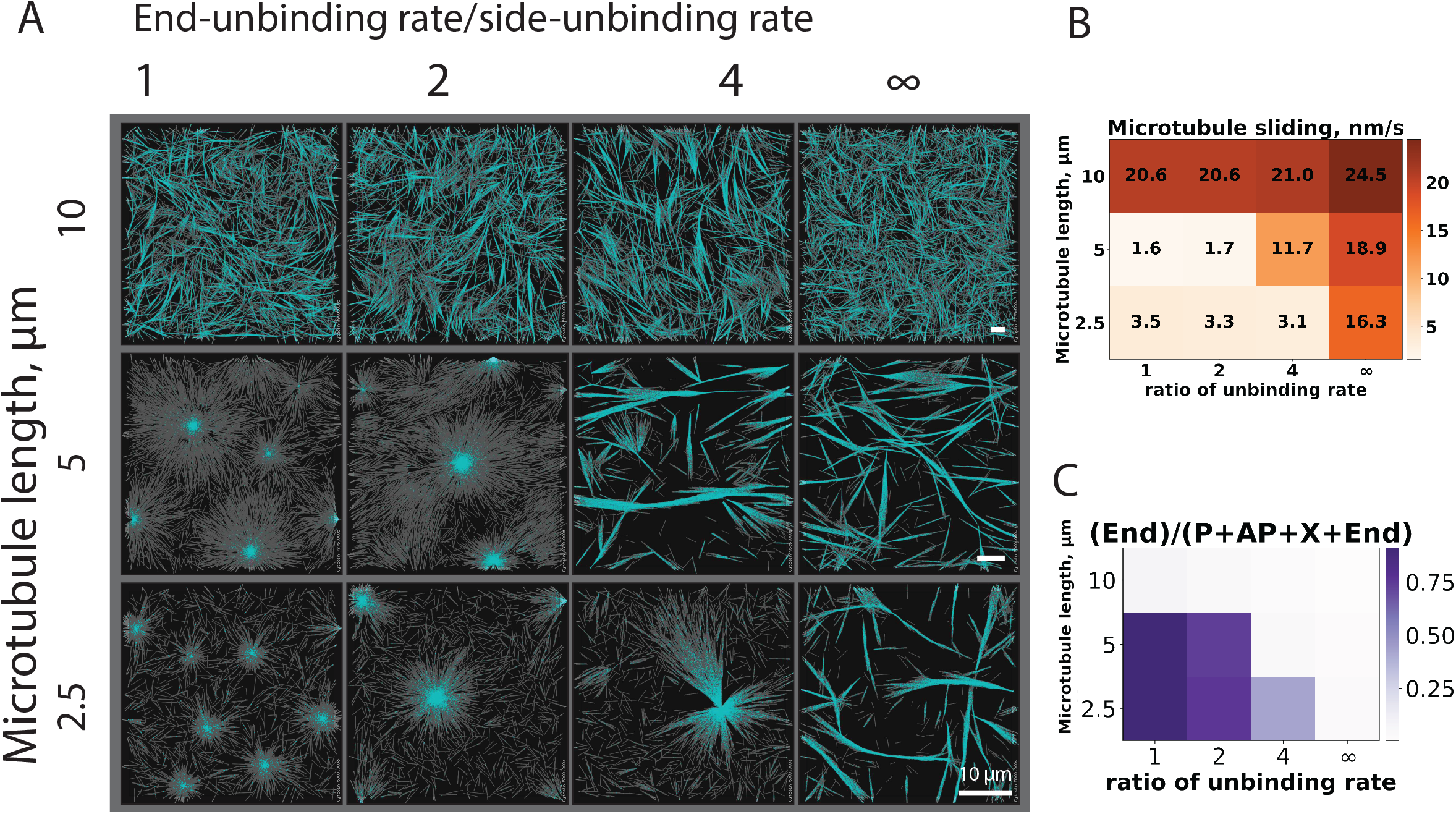
(A) Microtubule organizations with increasing motor end-unbinding rates for three different individual microtubule lengths at a constant total microtubule length per area of 5 μm/μm^2^. There are 16 KIF11 motors per microtubule. The simulation lasted for 60 min in a box of dimension: Lx μm × Ly μm × 0.2 μm, where Lx = Ly = 16 × (microtubule length). All scale bars are 10 μm. (B) Calculated mobility of microtubule minus ends in each simulation at 60 min. (C) Fraction of end links for each simulation shown in (A). See also Movie 4.

### Experimental demonstration of the existence of the ‘crawling mesh’ state

Finally, to test the prediction of the existence of ‘crawling mesh’ state made by our simulations, we performed experiments. Microtubules were nucleated from purified tubulin in a glass chamber and subsequently organized by purified KIF11 motor proteins (see Methods). We varied the tubulin concentration and saw that KIF11 forms contractile networks of asters with low densities of microtubules, and active nematic networks in higher densities, as shown previously (Fig. 7A, Movie 5) (Roostalu et al. 2018, Henkin et al. 2022). At the highest tubulin concentrations for a given amount of motor protein (where microtubule density and length are expected to be highest), the network instead formed a crosslinked mesh without obvious macroscopic ordering, corresponding to the ‘crawling mesh’ state found in the simulations. Incorporation of pre-polymerized, stabilized microtubule “seeds” with a spectrally separate fluorescent dye showed that despite the lack of macroscopic order, individual microtubules were highly motile, and translocated in the direction of their axis (Fig. 7B, Movie 6). This confirms experimentally the existence of the predicted ‘crawling mesh’, a new type of active microtubule network.

**Figure 7.**
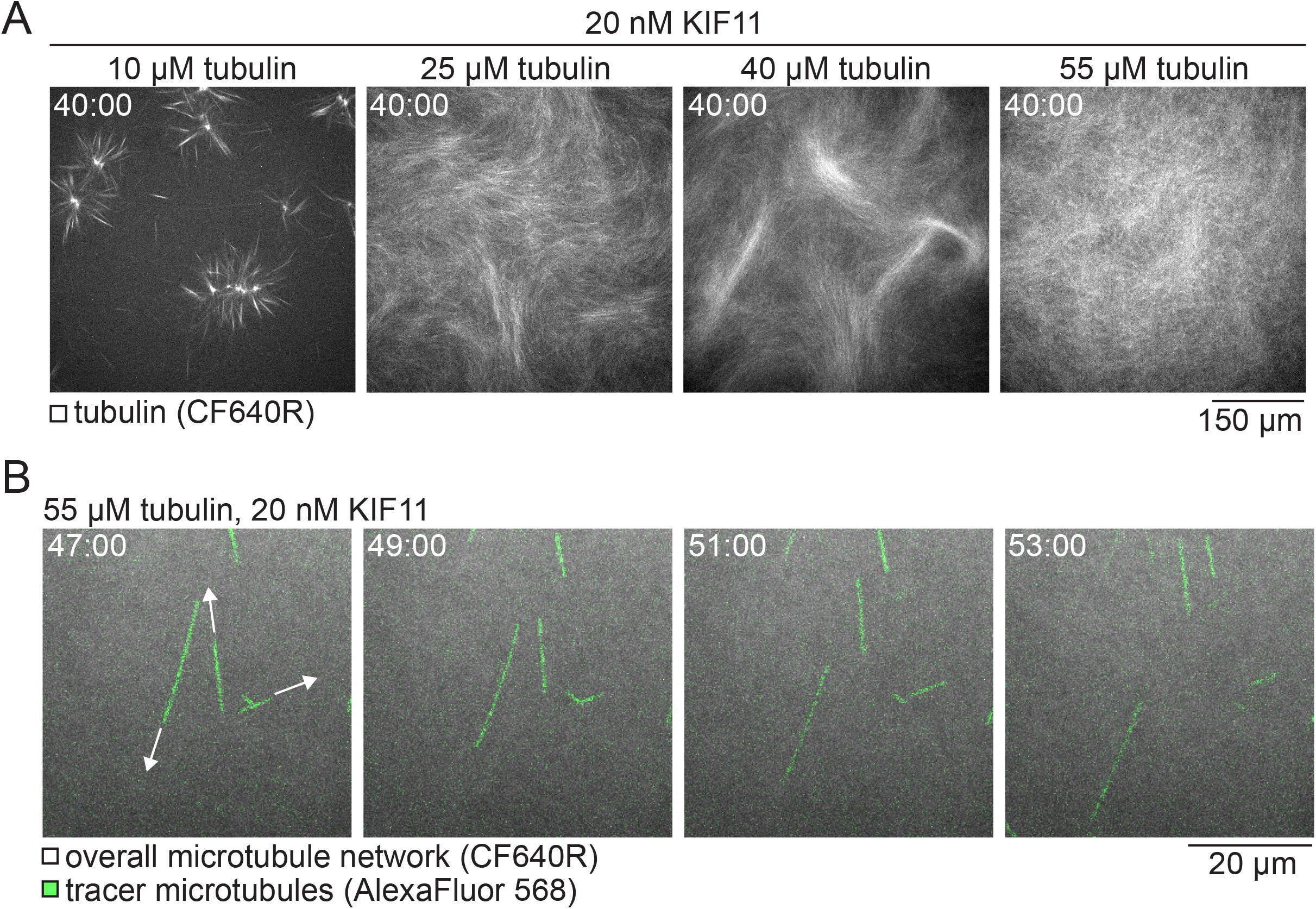
(A) Confocal fluorescence microscopy images showing CF640R-labeled tubulin in microtubule networks polymerized and organized in the presence of 20 nM KIF11. With increasing tubulin concentration, the macroscopic network state transitions from contractile (10 μM tubulin), to active nematic (25, 40 μM tubulin), to isotropic (55 μM tubulin). See also Movie 5. (B) Time course at higher magnification showing fluorescence of AlexaFluor568-labeled GMPCPP tracer microtubules (green) as they glide through an overall isotropic network organized by 20 nM KIF11 from 55 μM tubulin (CF640R-tubulin; gray). See also Movie 6.

## DISCUSSION

Microtubule/motor systems can self-organize into different types of active networks. Several important control parameters determining their organization, such as the densities of motors and microtubules or their speed of motion and growth, have been identified by experimental and theoretical studies (Roostalu et al. 2018, Rickman, Nedelec and Surrey 2019, Blackwell et al. 2016, Fürthauer et al. 2021). Here we focused on two important control parameters that so far escaped substantial theoretical investigation: (i) short-range attractive forces between microtubules, and (ii) the length of microtubules. In experiments, microtubule-microtubule attractive forces can be manipulated by adding crowding agents. Microtubule length is more difficult to control *in vitro*, and moreover can vary together with microtubule density and growth speed, making the influence of microtubule length on network organization harder to understand from experiments alone. Living cells, however, have evolved many mechanisms to control microtubule length, as this is clearly a key parameter controlling the organization of microtubule networks (Lacroix and Dumont 2022, Howard and Hyman 2007).

We were able to reproduce in our simulations of microtubule/motor networks the transition from contractile asters to networks of extensile bundles by increasing the strength of short range attraction between microtubules, as induced by crowding agents in previous experiments (Lemma et al. 2022). We found that the microtubule density needed to be high enough to allow re-mixing of microtubules between long-range extensile bundle connections, to prevent local microtubule polarity-sorting, as also observed in experiments (Lemma et al. 2022). A sufficiently high microtubule density together with a depletion force results in a turbulent state in which the microtubule bundles can extend continuously. At lower microtubule densities, bundles instead polarity-sort, because they fail to fuse and re-mix, explaining why a certain microtubule density is required to achieve a network of permanently extensile bundles (Sanchez et al. 2012, Lemma et al. 2022)

In experiments, control parameters are often interdependent, particularly when the same protein is involved in related processes. When microtubules polymerize in solution and the tubulin concentration is changed, this affects the number of microtubules, their growth speed and their length. Previous work showed that the transition from asters to nematic networks can be obtained with dynamic microtubules and motors in the absence of crowding agents when the tubulin concentration was increased (Roostalu et al. 2018). Simulations explained that the resulting increase in microtubule number and growth speed promoted nematic network formation by favouring microtubule side-to-side links over end links. The effect of the microtubule length remained however unexplored.

Here we find in our simulations that the formation of networks of extensile bundles is also promoted by increasing the microtubule length, essentially for the same reason, but beyond a certain threshold a qualitatively new state emerges: the ‘crawling mesh’ state. Long microtubules cannot easily reorient as the network is highly percolated. Instead, they continuously slide unidirectionally through the isotropic network. This state has probably been observed in previous experiments with high tubulin and high motor concentrations, but it has received less attention and was described as ‘unorganized’ or ‘stuck’ (Roostalu et al. 2018). Here we tested the prediction of the simulations experimentally and were indeed able to reproduce the transition from a contractile network forming asters to a network of extensile bundles and finally to the ‘crawling mesh’ state by increasing the tubulin concentration. Microtubules sliding through the ‘crawling mesh’ could be observed directly by labelling a subset of the microtubules.

It is known from past simulations that the microtubule end unbinding rate of the crosslinking motors needs to be slow enough to allow aster formation (Surrey et al. 2001, Roostalu et al. 2018, Rickman et al. 2019). We showed here that this control parameter shifts more generally the boundaries between the different network states. As aster formation becomes more difficult with an increasing end unbinding rate, the nematic network state becomes more accessible for short microtubules. Tuning this kinetic parameter in simulations can be useful from a practical point of view. Our simulations with the longer microtubules whose lengths can easily be reached in experiments are extremely time-consuming (2-3 weeks), because the simulation space had to be very large to avoid artifacts caused by the boundaries. The simulation space and time can however be reduced significantly using shorter microtubules and a higher end-unbinding rate to compensate for their stronger tendency to form contractile networks, while reproducing the same system tendencies and making experimentally testable predictions. This then allows one to simulate the experimentally observed network transitions with shorter microtubules (Henkin et al. 2022).

In conclusion, we have used computer simulations to show that the microtubule length and the strength of a short-range attraction between microtubules that mimics a crowder-induced depletion force are important control parameters for the active network organization of microtubule/motor systems. Our simulations are in good agreement with previous experiments and have made a prediction regarding the dynamic state of a highly percolated isotropic network that we could confirm experimentally. A strength of the simulations is that the different topologically defined types of motor crosslinks that characterize the network self-organizations can be extracted over the time course of its development, something that is less accessible in experiments and that can provide mechanistic insight into the principles that drive contractile versus extensile active network formation. Our results here expand our understanding of the effects that control parameters have on active microtubule networks and help to better understand the control of active network architectures in cells and may also help to engineer novel biomimetic or bioinspired materials.

## METHODS

### Model

We simulated active networks consisting of microtubules and motors using Cytosim. Our aim was to systematically explore the effects of key parameters and to monitor the system organization through a limited set of scalar quantities calculated automatically. The model is essentially as described earlier (Henkin et al. 2022, Roostalu et al. 2018, Rickman et al. 2019). In brief, microtubules are modelled as diffusing, flexible filaments repelling each other via soft-core interactions.

To study the influence of short-range attractive force between the microtubules induced by depletion force in the presence of crowders. We introduce a piecewise linear force that is repulsive below range *d*_0_, attractive between *d*_0_ and *d*_1_ and null above *d*_0_ + *d*_1_:

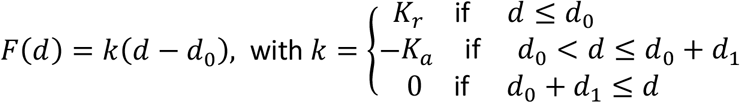

where *d* is the distance between two interacting vertices of the filaments. This force is characterized by distances *d_0_* and *d_1_*, and by *K_r_* and *K_a_* the repulsive and attractive force stiffnesses. In a bundle at equilibrium, the filaments are typically separated by *d_0_*, measured center-to-center. The force is orthogonal to the filament so as to permit sliding of the microtubules parallel to their axis.

The equilibrium *d_0_* is set to be 0.1 μm, *d_1_* is 0.32 μm and *K_r_* is set to be 50 pN/μm. We mimic the concentration dependent depletion strength by varying *K_a_* from 0.3 to 1.2 pN/μm, which corresponds to maximum force around 0.1 - 0.4 pN. In this way, the magnitude of the force is in the same order as the estimated depletion force between a pair of cytoskeletal filaments which is around 0.1-0.2 pN per binding site (Streichfuss et al. 2011, Hilitski et al. 2015).

Microtubules grow from a fixed number of nucleators by plus-end elongation. Microtubule crosslinking motors can bind stochastically to two microtubules at most and walk along them in the plus-end direction. The properties of the motor are set to mimic those of human kinesin-5 (KIF11, also known as Eg5 in Xenopus), as in previous work (Roostalu et al. 2018, Henkin et al. 2022). Microtubule bound motor can unbind with a higher rate at the end of microtubule than at the side. All simulations were performed in a flat and thin threedimensional geometry with periodic boundary conditions in the X and Y dimensions and reflecting boundaries in the much shorter Z dimension, allowing the formation of extended quasi-two-dimensional networks (Henkin et al. 2022, Roostalu et al. 2018, Rickman et al. 2019). The size of the simulation box is Lx = Ly = 16L where L is the microtubule length. The thickness of the box Lz is 0.2 μm in all simulations presented here, and we thus quantify the system’s density by the total length of microtubule divided by Lx × Ly. Detailed parameters of the model are available in Table S1.

### Nematic order parameter

In dimensionality *d*, the orientational order is characterized by a symmetric traceless *d*×*d* tensor (Gennes and Prost 1993, Doi 2013) built using the outer product ⊗:

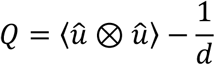

Where *û* is a unit *d*-vector directed along the axis of microtubules, and 〈·〉 denotes ensemble average over all microtubules. The scalar order parameter *S* is the largest eigenvalue of *Q*. Since our system is quasi-2D, we calculated a 2D order parameter, only considering the X and Y components of the filament’s 3D direction vectors, rescaled such that *û_x_*^2^ + *û_y_*^2^. By construction, *S* ∈ [0,1]. If the system remains isotropic (no alignment), *S* is close to 0. If the alignment is perfect (nematic or vectorial), *S* is equal to 1.

To capture the local nematic order, the sampling window must be smaller than the simulation box, but this window size must be chosen carefully. To differentiate aster from nematic bundle, we adjusted the window size to contain the largest aster observed, that is 10 μm × 10 μm. This lowers the order calculated for an aster since microtubules from the aster radiate in all directions. A large window size also lowers the order calculated when several bundles of different orientations are present in the window. Nevertheless, for sparse and thick bundles, the order value remains close to one in this work. To avoid overcounting the contribution of isolated microtubules, the order parameter of each sampling window is weighted by the number of microtubules in that window. The weighted average of the local order parameter of all windows gives the nematic order parameter of the entire system, referred hereon simply as *S*.

### Crosslink types

To characterize different types of connection made by motor between microtubules, we defined four types of crosslinks: P link, X link, AP link and end links as described before (Roostalu et al. 2018). P links connect parallel microtubules where the internal angle is smaller than 60 degrees; X links connect microtubules with an angle between 60 and 120 degrees. AP-links connect antiparallel microtubules with an angle between 120 and 180 degrees. End links connect one or both microtubule near their plus ends (within a distance of 10nm). We use the proportion of end links out of all other links, (end links/all links), to quantify the degree of microtubules end clustering (aster). Among the non-end links (P+AP+X links), the relative proportion of P link, AP link and X link are also monitored.

### Microtubule mobility

To calculate the overall speed of microtubule motion, we extracted the positions of microtubule minus ends at regular time intervals Δt. We then calculated the displacement component parallel to the microtubule axis. Averaging this signed scalar displacement for all microtubules and dividing by Δt gave the overall averaged microtubule speed. A large Δt = 200 s was chosen to ensure that the contribution of diffusion becomes negligible relative to the active motion generated by the motors. As a matter of convention, a positive speed indicates minus-end leading sliding (as driven by plus-end directed motors).

### Principal component and clustering analysis

We aim to categorize the simulated microtubule organization based on the six descriptors of the network state: local order parameter, microtubule mobility, P links, AP links, X links, and end links using clustering algorithm. To reduce the dimension of data set, we perform principal component analysis (PCA) (Pearson 1901) on the six descriptors using the Scikit-learn library (https://scikit-learn.org). In our analysis, the first two principal components can explain 94% of the variances where the loading vectors are tabulated in Fig S1A and Fig. S1C. We then categorize the microtubule network in this subspace using K-means clustering method(Lloyd 1982) to obtain the distinct clusters.

### Experimental self-organization assay

Samples for the self-organization assays in Fig. 7 were prepared similarly to previous work (Henkin et al. 2022). Pig brain tubulin and recombinant KIF11-mGFP were prepared as previously described (Henkin et al. 2022, Consolati et al. 2022, Roostalu et al. 2018). Passivated glass coverslips were prepared as described (Consolati et al. 2022), however glass was cleaned by sonication in acetone followed by plasma cleaning in place of sonication in piranha solution. Chambers were prepared using two layers of 10 μm thick double-stick adhesive tape (Nitto Denko) for a final chamber height of about 20 μm. Tubulin (including CF640R-labeled tubulin at a final label ratio of 3.5%) and KIF11-mGFP (for reported concentrations referring to monomers) were mixed into an assay buffer on ice, spun at 13.3k rpm at 4° C in a table-top centrifuge, recovering the supernatant. The supernatant was transferred to a tube at room temperature, and mixed with BRB80 (80 mM PIPES, 1 mM MgCl_2_, 1 mM EGTA, pH 6.8) or GMPCPP seeds (AlexaFluor568; 14% labeling ratio) diluted in BRB80, for final assay component concentrations including 0.68 mg/mL glucose oxidase, 0.17 mg/mL catalase, 0.9 mg/mL β-casein, in a buffer of 40 mM PIPES, 1 mM EGTA, 1.6 mM MgCl_2_, 0.9 mM ATP, 0.58 mM GTP, 32 mM glucose, 3.2 mM β-mercaptoethanol, and 1 μM docetaxel, for a final pH of 6.9 – 6.95. Chambers were preheated to 33° C on a heat block and washed with BRB80 buffer just before loading the final sample and sealing with silicone vacuum grease. Imaging was performed on a spinning disk confocal microscope at 33° C around 3 minutes after the initial temperature shift, which stimulates microtubule nucleation and growth. Timestamps refer to time since the beginning of imaging. Panels in Fig. 7A are single slices from the chamber midplane, whereas panels in Fig. 7B are maximum-intensity projections of 5 slices, 1 μm apart, around the chamber midplane. Intensities are adjusted independently for each experiment.

## ACKNOWLEDGMENTS

We thank the members of the Surrey lab for useful discussions. Simulations were performed on the high-performance computing cluster at the Centre for Genomic Regulation (CRG) in Barcelona, Spain. This work was supported by the Spanish Ministry of Economy, Industry and Competitiveness to the CRG-EMBL partnership, the Centro de Excelencia Severo Ochoa and the CERCA Programme of the Generalitat de Catalunya. W.-X.C. is supported by a Human Frontier Science Program fellowship (HFSP LT000682/2020-C). F.N. is supported by the Gatsby Charitable Foundation (Grant PTAG-024). F.N. and T.S. acknowledge support from the European Research Council (ERC Synergy Grant, Project 951430).

## AUTHOR CONTRIBUTIONS

W.-X.C., G.H., F.N., and T.S. designed research; W.-X.C. performed theoretical research and G.H. experimental research; W.-X.C., and F.N. analyzed data; and W.-X.C., G.H., F.N., and T.S. wrote the paper.

## FIGURE LEGENDS

**Figure S1.**
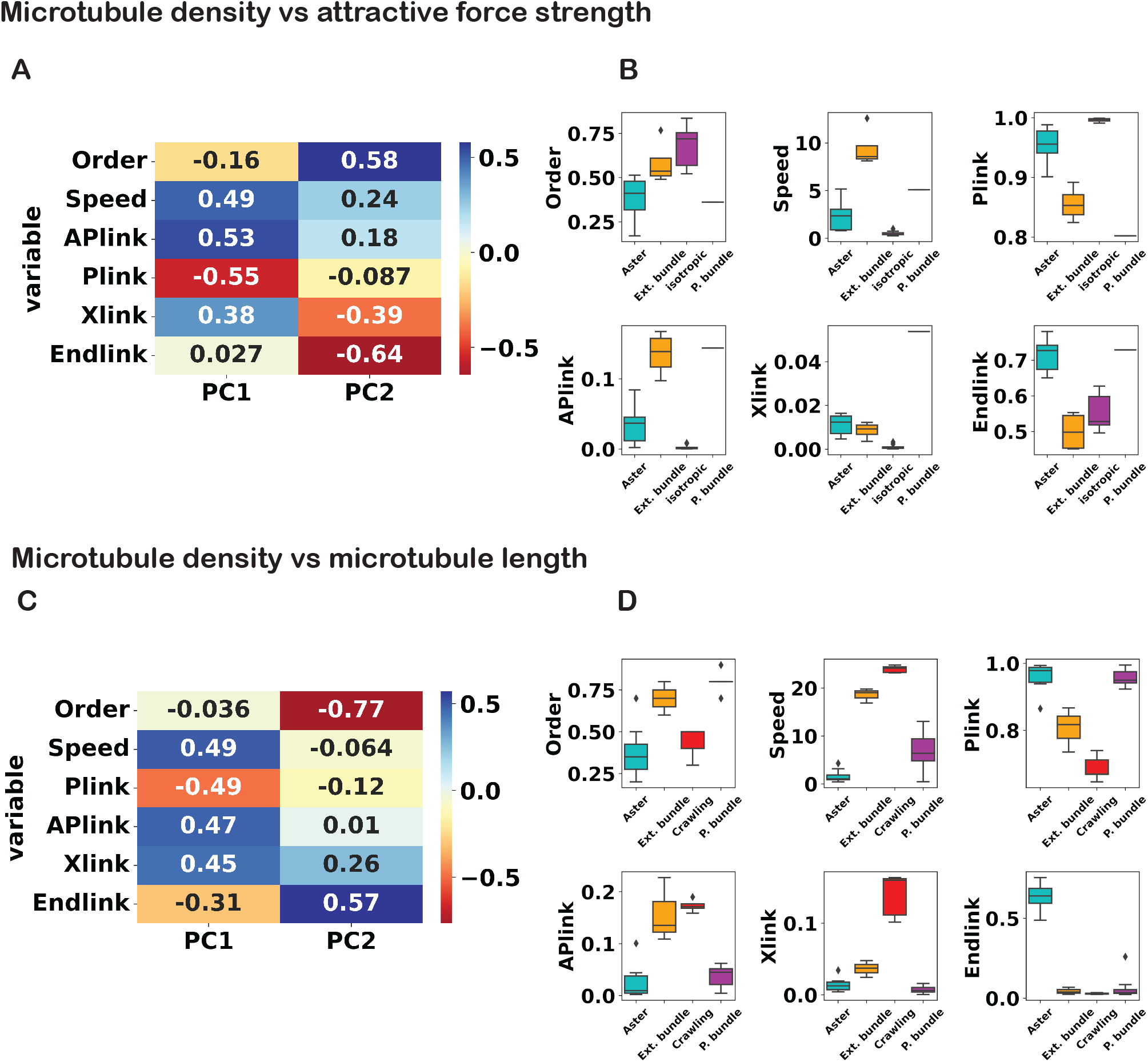
Breakdown of the loading vectors of each feature for the two principal components corresponding to analysis in (A) Fig. 3B and (C) Fig. 5B. Boxplot of each feature grouped by clusters corresponding to analysis in (B) Fig. 3B and (D) Fig. 5B.

**Figure S2.**
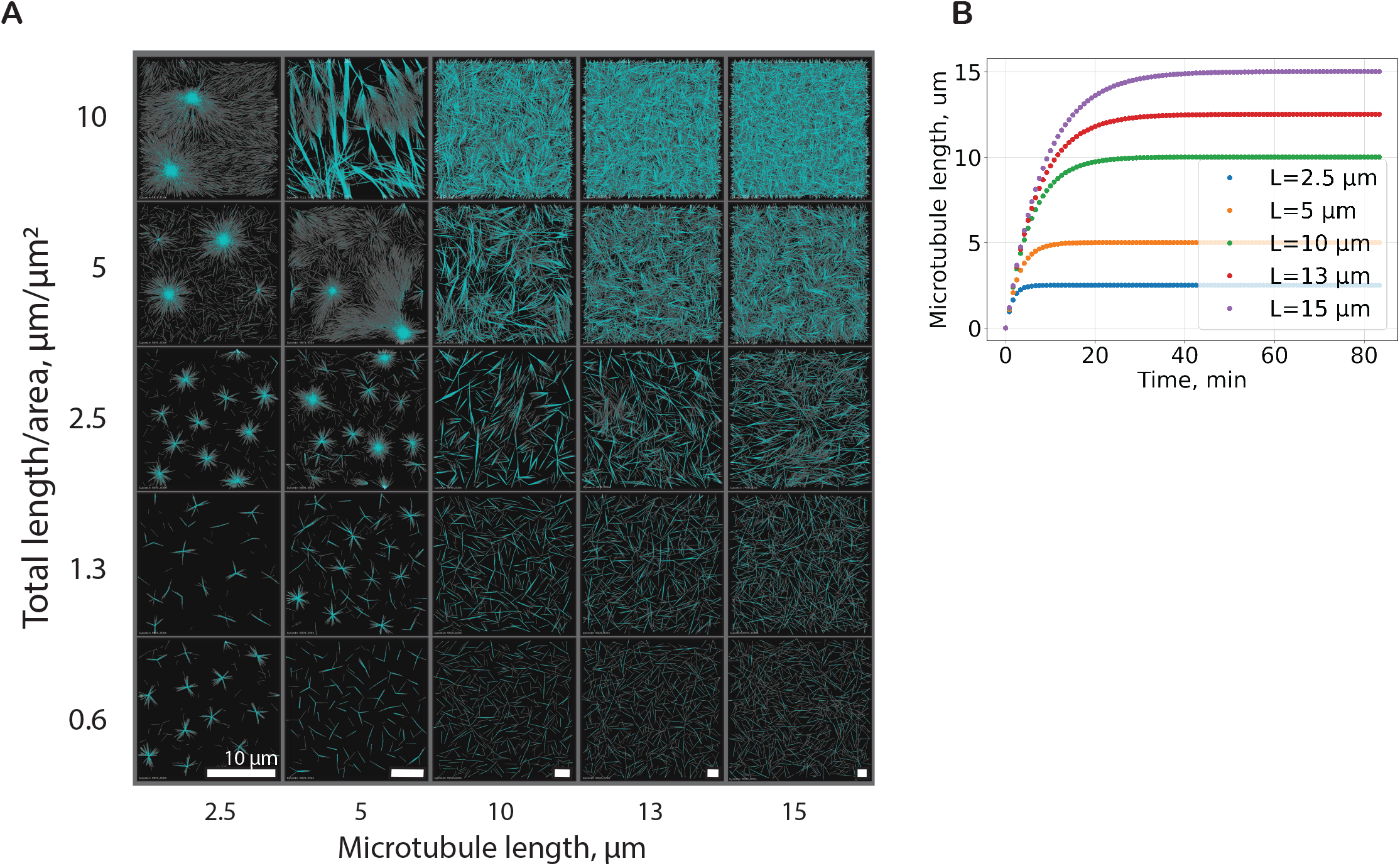
(A) Organizational phase space of the microtubule network at varying microtubule densities and lengths. The KIF11 motor to microtubule ratio is 16. The simulation was simulated for 60 min in a box of dimensions Lx μm × Ly μm × 0.2 μm, where Lx = Ly = 16 × (microtubule length). The full view of the simulation box is shown here. All scale bars are 10 μm. (B) Time course of the microtubule mean lengths for different maximum lengths.

**Figure S3.**
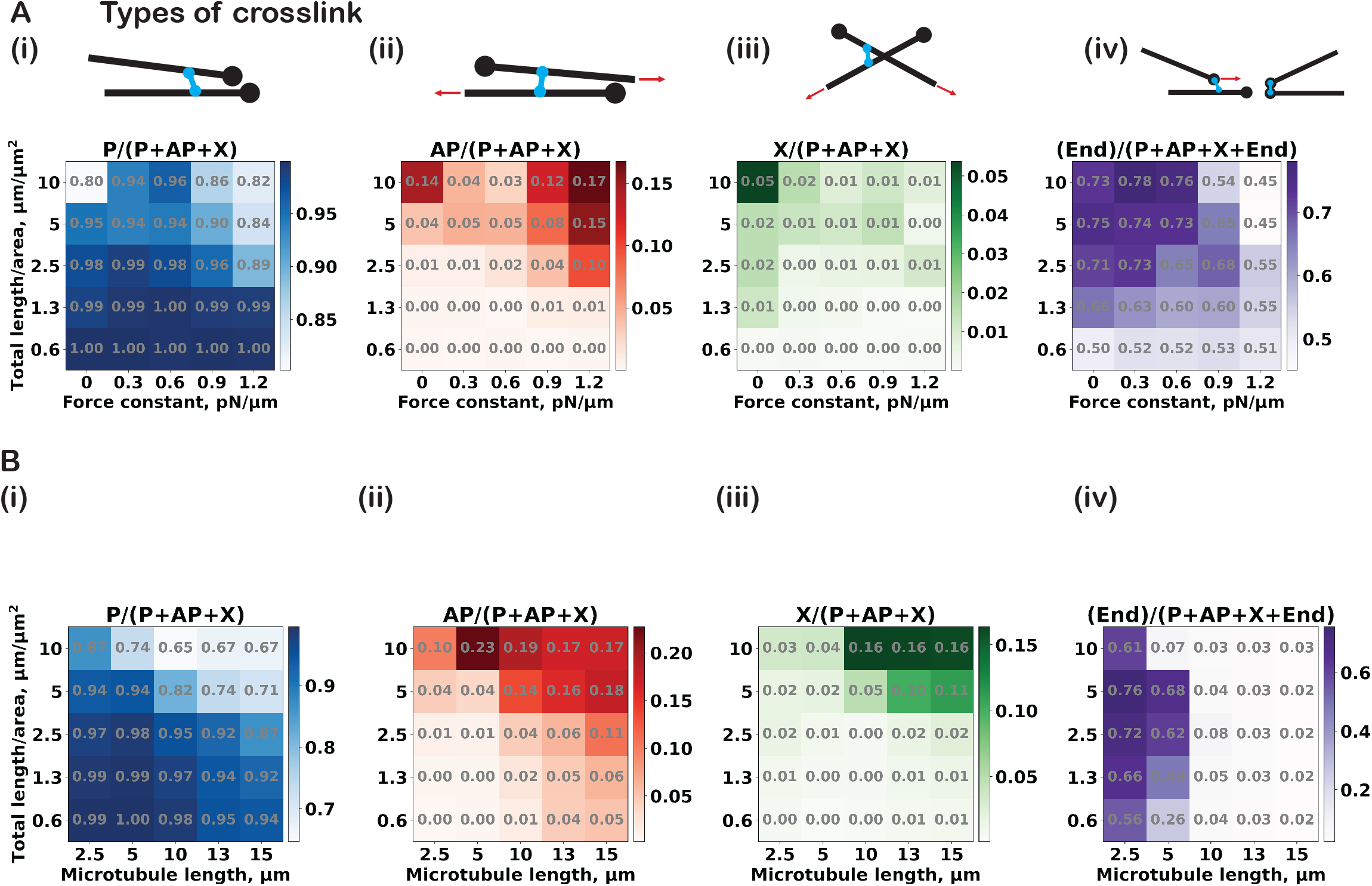
Numerical version of (A) Fig. 3A and (B) Fig. 5A.

## MOVIE LEGENDS

**Movie 1.** Time course of simulated self-organizing networks driven by a KIF11-like motor (cyan) without attractive bundling force that form small asters (top left, density: 1.3 μm/μm^2^, k_a_: 0 pN/μm), large asters (top right, density: 5 μm/μm^2^, k_a_: 0 pN/μm), and with attractive bundling force that form parallel bundles (bottom left, density: 1.3 μm/μm^2^, k_a_: 1.2 pN/μm), and extensile bundles (bottom right, density: 5 μm/μm^2^, k_a_: 1.2 pN/μm)

**Movie 2.** Time course of the organization of microtubules (gray) and KIF11 motors (cyan) in the absence of an attractive bundling force that form asters (left, L: 2.5 μm), parallel bundles (middle, L: 5 μm), and a network of extensile bundles (right, L: 15 μm). Total microtubule length per area is 2.5 μm/μm^2^.

**Movie 3.** (A) Time course of the organization of microtubules (gray) and KIF11 motors (cyan) in the absence of an attractive bundling force that form asters (left, L: 2.5 μm), extensile bundles (middle, L: 5 μm) and a ‘crawling mesh’ organization (right, L: 15 μm). (B) Time course of the ‘crawling mesh’ network with all microtubules shown in grey filaments (left) and a subset of microtubules colored distinctly (right). Total microtubule length per area is 10 μm/μm^2^.

**Movie 4.** Time course of the organization of microtubules (gray) and KIF11 motors (cyan) in the absence of an attractive bundling force that form asters at low end-unbinding rate (left, density: 5 μm/μm^2^, L: 5 μm; ratio of end-unbinding rate to side-unbinding rate: 2) and extensile bundles (right, density: 5 μm/μm^2^, L: 5 μm; ratio of end-unbinding rate to side-unbinding rate: infinite).

**Movie 5.** Time-course of experimental self-organizing microtubule networks in 20 μm thick chambers, driven by 20 nM KIF11, in the presence of 10, 25, 40 and 55 μM tubulin (CF640R-tubulin; final labeling ratio 3.5%); confocal images for each experiment are taken at the chamber midplane.

**Movie 6.** Translocation of pre-polymerized, stable GMPCPP tracer microtubules (14% AlexaFluor 568; left) in an overall isotropic network driven by 20 nM KIF11 (55 μM tubulin, 3.5% CF640R-tubulin; right) in a 20 μm thick chamber, demonstrating the ‘crawling mesh’ state.

**Table S1.**
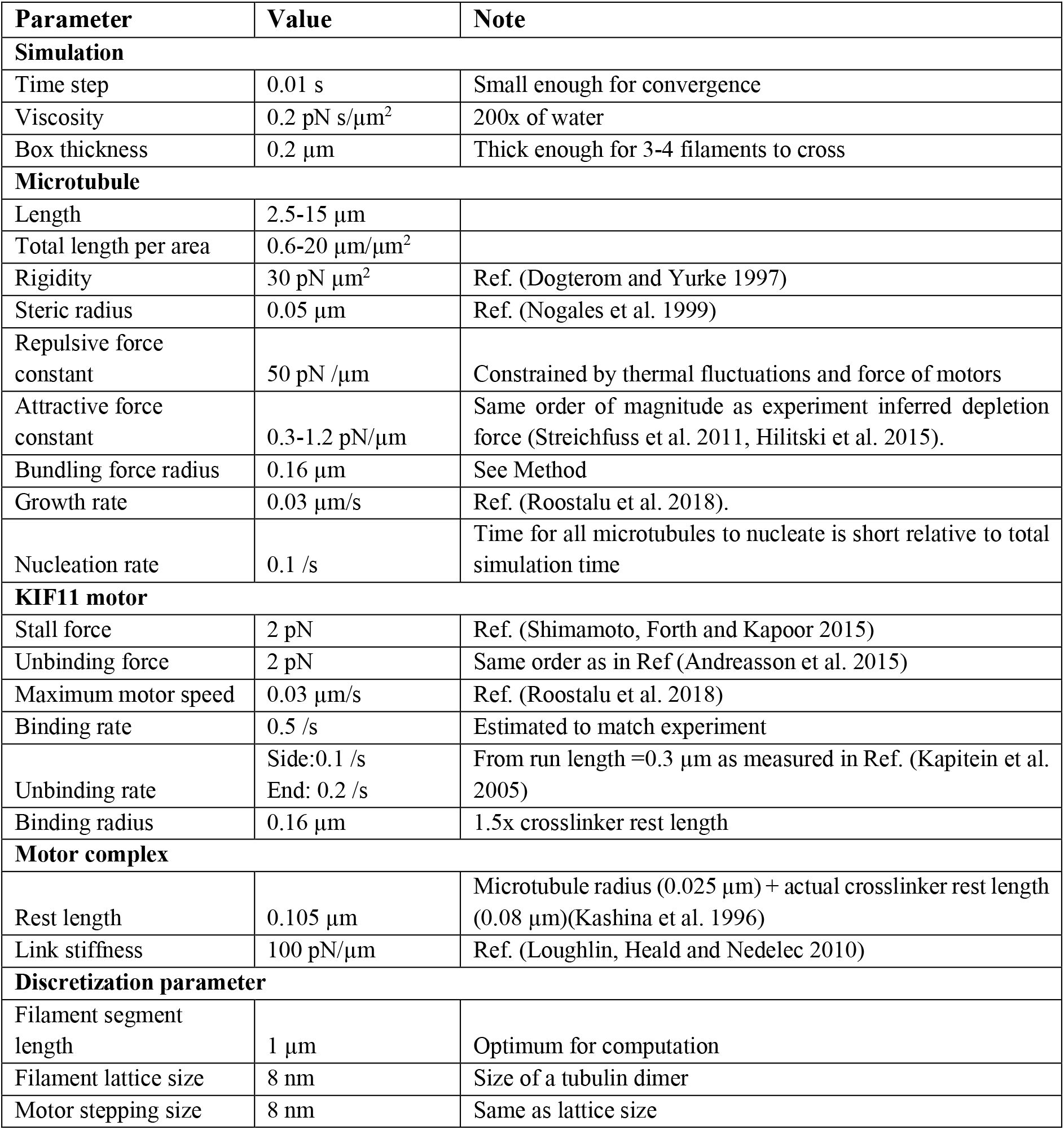

| Parameter | Value | Note |
| --- | --- | --- |
| **Simulation** |  |  |
| Time step | 0.01 s | Small enough for convergence |
| Viscosity | 0.2 pN s/$\mu\text{m}^2$ | 200x of water |
| Box thickness | 0.2 $\mu\text{m}$ | Thick enough for 3-4 filaments to cross |
| **Microtubule** |  |  |
| Length | 2.5-15 $\mu\text{m}$ | |
| Total length per area | 0.6-20 $\mu\text{m}/\mu\text{m}^2$ | |
| Rigidity | 30 pN $\mu\text{m}^2$ | Ref. (Dogterom and Yurke 1997) |
| Steric radius | 0.05 $\mu\text{m}$ | Ref. (Nogales et al. 1999) |
| Repulsive force constant | 50 pN /$\mu\text{m}$ | Constrained by thermal fluctuations and force of motors |
| Attractive force constant | 0.3-1.2 pN/$\mu\text{m}$ | Same order of magnitude as experiment inferred depletion force (Streichfuss et al. 2011, Hilitski et al. 2015). |
| Bundling force radius | 0.16 $\mu\text{m}$ | See Method |
| Growth rate | 0.03 $\mu\text{m}/\text{s}$ | Ref. (Roostalu et al. 2018). |
| Nucleation rate | 0.1 /s | Time for all microtubules to nucleate is short relative to total simulation time |
| **KIF11 motor** |  |  |
| Stall force | 2 pN | Ref. (Shimamoto, Forth and Kapoor 2015) |
| Unbinding force | 2 pN | Same order as in Ref (Andreasson et al. 2015) |
| Maximum motor speed | 0.03 $\mu\text{m}/\text{s}$ | Ref. (Roostalu et al. 2018) |
| Binding rate | 0.5 /s | Estimated to match experiment |
| Unbinding rate | Side:0.1 /s<br>End: 0.2 /s | From run length =0.3 $\mu\text{m}$ as measured in Ref. (Kapitein et al. 2005) |
| Binding radius | 0.16 $\mu\text{m}$ | 1.5x crosslinker rest length |
| **Motor complex** |  |  |
| Rest length | 0.105 $\mu\text{m}$ | Microtubule radius (0.025 $\mu\text{m}$) + actual crosslinker rest length (0.08 $\mu\text{m}$)(Kashina et al. 1996) |
| Link stiffness | 100 pN/$\mu\text{m}$ | Ref. (Loughlin, Heald and Nedelec 2010) |
| **Discretization parameter** |  |  |
| Filament segment length | 1 $\mu\text{m}$ | Optimum for computation |
| Filament lattice size | 8 nm | Size of a tubulin dimer |
| Motor stepping size | 8 nm | Same as lattice size |

